# Meta-analysis of RNA Sequencing Data of Arabidopsis and Rice under Hypoxia

**DOI:** 10.1101/2022.06.23.497423

**Authors:** Keita Tamura, Hidemasa Bono

## Abstract

Hypoxia is an abiotic stress in plants. Flooding resulting from climate change is a major crop threat that increases the risk of hypoxic stress. The molecular mechanisms underlying hypoxia in plants have been elucidated in recent years, but new genes related to this stress remain to be discovered. Thus, we aimed to perform a meta-analysis of RNA sequencing (RNA-Seq) data of Arabidopsis (*Arabidopsis thaliana*) and rice (*Oryza sativa*) under hypoxia. We collected 29 (Arabidopsis) and 26 (rice) pairs of RNA-Seq data involving hypoxic (including submergence) and normoxic (control) treatments and extracted the genes that were commonly upregulated or downregulated in the majority of the experiments. The meta-analysis revealed 40 and 19 commonly upregulated and downregulated genes, respectively, in the two species. Several WRKY transcription factors and cinnamate-4-hydroxylase were commonly upregulated, but their involvement in hypoxia remains unclear. Our meta-analysis identified candidate genes for novel molecular mechanisms in plants under hypoxia.

## 1. Introduction

Flooding is a major abiotic stress for crop production, and water availability extremes (drought and flooding) have become frequent, owing to climate change [1]. Flooding can be classified into waterlogging (partial coverage of water) and submergence (complete coverage of water), both of which limit oxygen availability to plants and lead to hypoxia [2]. Considering the importance of molecular oxygen in eukaryotes, several studies have elucidated and compared the molecular mechanisms by which plants and animals respond to hypoxia [3,4]. In metazoans, hypoxia-inducible factor (HIF) is a transcription factor (TF) that regulates the transcriptional changes induced by oxygen availability [3,5]. In plants, group VII ethylene response factor TFs (ERFVIIs) induce transcriptional changes under hypoxia [3]. HIF and ERFVIIs are a functionally analogous pair that stabilizes under hypoxia and degrades under oxygenated conditions; however, their pathways are mechanistically different [3].

The molecular mechanisms of plants under hypoxic conditions, including flooding, submergence, and waterlogging, have been extensively studied in Arabidopsis (*Arabidopsis thaliana*) and rice (*Oryza sativa*) [2]. Ethylene is entrapped in plants upon flooding, causing signaling cascades for adaptation to hypoxia [6]. The discovery of ERFVII genes, such as *SUBMERGENCE 1A* (*SUB1A*) [7,8], *SNORKEL1* (*SK1*), and *SNORKEL2* (*SK2*) [9], in rice varieties with submergence tolerance has led to the study of this gene family in Arabidopsis [6]. In Arabidopsis, five ERFVII TFs have been identified [10,11]. Among them, *HYPOXIA RESPONSIVE ERF1* (*HRE1*) and *HRE2* are expressed under hypoxia [12], whereas *RELATED TO APETALA2*.*12* (*RAP2*.*12*), *RAP2*.*2*, and *RAP2*.*3* are constitutively expressed under normoxia [11]. Arabidopsis has 49 core hypoxia-responsive genes (HRGs), including *HRE1* and *HRE2* [13]. Stabilized ERFVII TFs under hypoxia are involved in the upregulation of over half of the 49 HRGs [14]. Transactivation studies of Arabidopsis ERFVII TFs indicated that *RAP2*.*12, RAP2*.*2*, and *RAP2*.*3* act redundant to activate HRGs, whereas *HRE1* and *HRE2* play minor roles as activators [11]. The stability of ERFVII TFs with oxygen-sensing mechanisms or protein–protein interactions with ERFVII TFs have been extensively studied [2,6]. However, other molecular mechanisms outside the control of ERFVII TFs remain to be determined.

Identifying novel molecular mechanisms under specific biological conditions is an important biological study, but hypothesis construction is generally influenced by well-studied theories and publication bias. Thus, a meta-analysis, which is a data-driven and unbiased collective analysis of multiple studies, is a promising strategy to provide novel insights [15,16]. Recently, the number of transcriptome analyses has increased dramatically, and most transcriptome studies have performed RNA sequencing (RNA-Seq) using the Illumina platform [All of Gene Expression (AOE); https://aoe.dbcls.jp] [17]. This abundance of studies makes a meta-analysis of RNA-Seq data feasible using publicly available data. This strategy can lead to the identification of novel hypoxia-inducible genes in human cell lines or tissue specimens [15].

In this study, we aimed to perform a meta-analysis of public RNA-Seq data of Arabidopsis and rice under hypoxia to identify novel hypoxia-inducible genes in plants. Experiments involving hypoxic treatments, including submergence and waterlogging, showed significant enrichment of genes with hypoxia-related gene ontology (GO) in the collection of frequently upregulated genes under this stress. The expression of several WRKY TFs and cinnamate-4-hydroxylase (C4H) was commonly upregulated. Our datasets contribute to the functional characterization of novel hypoxia-inducible genes in plants.

## 2. Materials and Methods

### 2.1. Curation of Public Gene Expression Data

To obtain public gene expression data related to hypoxia responses in plants, we searched Gene Expression Omnibus (GEO) [18] using the following keywords: (((“hypoxia” OR “hypoxic” OR “low oxygen” OR “submergence” OR “submerged” OR “waterlogging”) AND “Arabidopsis thaliana”[porgn:__txid3702]) AND “gse”[Filter] AND “Expression profiling by high throughput sequencing”[Filter]) for Arabidopsis, and (((“hypoxia” OR “hypoxic” OR “low oxygen” OR “submergence” OR “submerged” OR “waterlogging”) AND “Oryza sativa”[porgn:__txid4530]) AND “gse”[Filter] AND “Expression profiling by high throughput sequencing”[Filter]) for rice. Using the list of GSE accession numbers matched with the keywords, we obtained the corresponding SRP accession numbers and metadata with the Python package pysradb (v1.1.0). The obtained metadata were manually curated to focus on RNA-Seq of total mRNA and paired experiments of hypoxic and normoxic treatments. As a result, 29 (Arabidopsis) and 26 (rice) pairs of RNA-Seq data involving hypoxic (including submergence and waterlogging) and normoxic (control) treatments were created for this meta-analysis. The list of the RNA-Seq data used for this meta-analysis is available at figshare files 1a,b [19].

### 2.2. Gene Expression Quantification

FASTQ-formatted files for each RNA-Seq run accession number were retrieved using the SRA Toolkit (v2.11.0) (https://github.com/ncbi/sra-tools) with prefetch and fasterq-dump commands. FASTQ-formatted files for the same experiment were concatenated. Quality control of the raw reads was performed using Trim Galore (v0.6.7) (https://github.com/FelixKrueger/TrimGalore) with default settings. Transcript quantification was conducted using Salmon (v1.6.0) [20] against reference cDNA sequences downloaded from Ensembl Plants [21] (Arabidopsis_thaliana.TAIR10.cdna.all.fa.gz for Arabidopsis and Oryza_sativa.IRGSP-1.0. cdna.all.fa.gz for rice) with default settings (“-l A” was specified for automatic library type detection). These processes were performed using a set of scripts available at https://github.com/bonohu/SAQE [22] with small modifications. Quantitative RNA-Seq data, which were calculated as transcripts per million (TPM), are available at figshare (figshare files 2a,b) [19].

### 2.3. Calculation of HN-Ratio and HN-Score

To normalize the gene expression between different studies, the change in gene expression levels between each hypoxia and normoxia pair was measured as the HN-ratio [15]. The HN-ratio *R* was calculated using the following equation:

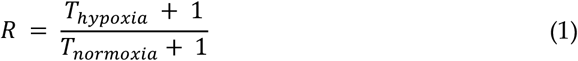

where *T*_*hypoxia*_ and *T*_*normoxia*_ are the TPM values of each gene in the hypoxic and normoxic treatments, respectively.

The HN-score [15] of each transcript was calculated to evaluate the changes in gene expression under hypoxia across experiments. We treated biological replicates in the same condition as individual experiment. In this study, transcripts with an HN-ratio of 2 or greater were considered to be upregulated, whereas those with an HN-ratio of 0.5 or less were considered to be downregulated. The HN-score of each transcript was calculated by subtracting the number of experiments with downregulated expression from the number of experiments with upregulated expression. The HN-ratio and HN-score were calculated using a set of scripts on https://github.com/no85j/hypoxia_code [15]. The HN-ratio and HN-score are available at figshare (figshare files 3a,b, and 4a,b) [19].

### 2.4. Functional Annotation of Transcripts

For the Arabidopsis data, the “Gene name” and “Gene description” for each transcript were obtained from Ensembl Biomart (Arabidopsis thaliana genes (TAIR10)) [23]. To find orthologs in human, we searched each transcript against human proteins (Homo_sapiens.GRCh38.pep.all.fa.gz) with an E-value cutoff of 1e-5 by blastx (v2.12.0) program and obtained the “Gene name” and “Gene description” (Human (GRCh38.p13)) for the top hits.

For the rice data, the “Gene name” and “Gene description” for each transcript were obtained from Ensembl Biomart (Oryza sativa Japonica Group genes (IRGSP-1.0)). To find orthologs in Arabidopsis, we searched each transcript against Arabidopsis proteins (Arabidopsis_thaliana.TAIR10.pep.all.fa.gz) with an E-value cutoff of 1e-10 by blastx (v2.12.0) and obtained the “Gene name” and “Gene description” (Arabidopsis thaliana genes (TAIR10)) for the top hits. Similarly, orthologs in humans were obtained as described in the Arabidopsis data.

### 2.5. Gene Set Enrichment Analysis

Gene set enrichment analysis was performed using Metascape (https://metascape.org/) [24] with the default settings (enrichment analysis with GO Biological Processes, KEGG Pathway and WikiPathways; p-value < 0.01, a minimum count of 3, and an enrichment factor > 1.5 (the enrichment factor is the ratio between the observed counts and the counts expected by chance)), or modified the ontology sources to GO Molecular Functions or GO Cellular Components. All genes in the genome have been used as the enrichment background, and up to the top 20 enriched clusters have been visualized. Rice transcripts were analyzed using orthologous transcript IDs in Arabidopsis. Venn diagrams were constructed using a web tool (https://bioinformatics.psb.ugent.be/webtools/Venn/).

### 2.6. Protein–Protein Interaction Analysis

Protein–Protein Interaction was analyzed and visualized using STRING (v11.5) (https://string-db.org/) [25] with default settings.

## 3. Results

### 3.1. Curation of RNA-Seq Data for Meta-analysis

Considering that hypoxia in plants has been associated with submergence and waterlogging, we used “submergence,” “waterlogging,” and “hypoxia” as keywords in the GEO search. The searches were refined in RNA-Seq data because unlike microarray data with various platforms, most RNA-Seq data are based on the Illumina platform, rendering them suitable for comparative analyses between studies. We collected 29 (Arabidopsis) and 26 (rice) pairs of RNA-Seq data of hypoxic and normoxic treatments (figshare files 1a,b) [19]. The data for Arabidopsis included hypoxic and submergence treatments. Although experiments with some mutants or transgenic lines were included, all data for Arabidopsis were derived with Col-0 background plants. 21 of the 29 pairs of the RNA-Seq data were sampled from seedlings, and the rest eight data were from leaves (figshare file 1a) [19]. The rice data included submergence or waterlogging treatments. M202(Sub1) [8], Varshadhan, and Rashpanjor [26] are flood-tolerant cultivars, whereas the majority of the experiments used flood-intolerant cultivars. 12 of the 26 pairs of the RNA-Seq data were sampled from above ground parts, and the rest 14 data were from roots (figshare file 1b) [19].

### 3.2. Characteristics of Upregulated Transcripts under Hypoxia

The meta-analysis of the RNA-Seq data of Arabidopsis and rice under hypoxia was performed by calculating the HN-score and HN-ratio of all transcripts [15]. The HN-score of each transcript is the difference between the number of experiments with downregulated expression and the number of experiments with upregulated expression; therefore, higher scores indicate global trends of upregulation across the experiments, whereas lower (minus) scores indicate global trends of downregulation across the experiments. We defined the top 1% transcripts with the highest and lowest HN-scores in the meta-analysis as upregulated and downregulated, respectively (figshare files 5a–d) [19]. The HN-score ranges of the upregulated and downregulated transcripts are summarized in Table 1.

**Table 1.** HN-scores of hypoxia-inducible transcripts identified in the meta-analysis.

| Species | Up or Down | HN-score (S) | No. of transcripts |
| --- | --- | --- | --- |
| Arabidopsis | upregulated | $11 \leq S \leq 26$ | 561 |
| Arabidopsis | downregulated | $-21 \leq S \leq -9$ | 493 |
| Rice | upregulated | $11 \leq S \leq 21$ | 606 (477)<sup>1</sup> |
| Rice | downregulated | $-22 \leq S \leq -10$ | 570 (489)<sup>1</sup> |

<sup>1</sup> Numbers in the parentheses are the number of transcripts annotated to Arabidopsis orthologs.

Gene set enrichment analysis of the upregulated transcripts in Arabidopsis and rice showed the enrichment of hypoxia-related terms in both species (Figure 1a,b). In Arabidopsis, the most significantly enriched term was GO:0071456 (cellular response to hypoxia) (Figure 1a). This was also the same when we analyzed hypoxic treatments and submergence treatments separately (figshare file 7) [19]. The highest HN-score of 26 was observed in AT4G24110.1, AT5G47910.1 (*RBOHD*), and AT1G76650.3 (*CML38*), all of which were included in the term GO:0071456 (figshare file 5a) [19]. AT4G24110, known as *HYPOXIA-RESPONSIVE UNKNOWN PROTEIN 40* (*HUP40*), is a member of 16 HUP genes in Arabidopsis [27]. RESPIRATORY BURST OXIDASE HOMOLOGUE D (RBOHD) is a NADPH oxidase involved in hydrogen peroxide (H_2_O_2_) production under hypoxia [28] and in plant immunity [29]. CALMODULIN-LIKE 38 (CML38) is a core hypoxia response calcium sensor protein [30]. In rice, the most significantly enriched term was GO:0001666 (response to hypoxia), the parent term of GO:0071456 (https://www.ebi.ac.uk/QuickGO/term/GO:0001666). The highest HN-score of 21 was observed in Os06t0605900-01 (*OsFbox316*) and Os01t0129600-00, of which only Os01t0129600-00 was included in the term GO:0001666 (figshare file 5b) [19]. The OsFbox316 ortholog in maize is upregulated under waterlogging [31]. Os01g0129600 (Os01t0129600-00) is similar to LOB DOMAIN-CONTAINING PROTEIN 40 (LBD40). Arabidopsis *LBD40* expression, together with *LBD4* and *LBD41* expression, is upregulated under hypoxia [32]. We also performed the gene set enrichment analysis against GO Molecular Function and GO Cellular Component (figshare file 8) [19]. Unlike GO Biological Process, the most significantly enriched term did not match between Arabidopsis and rice.

**Figure 1.**
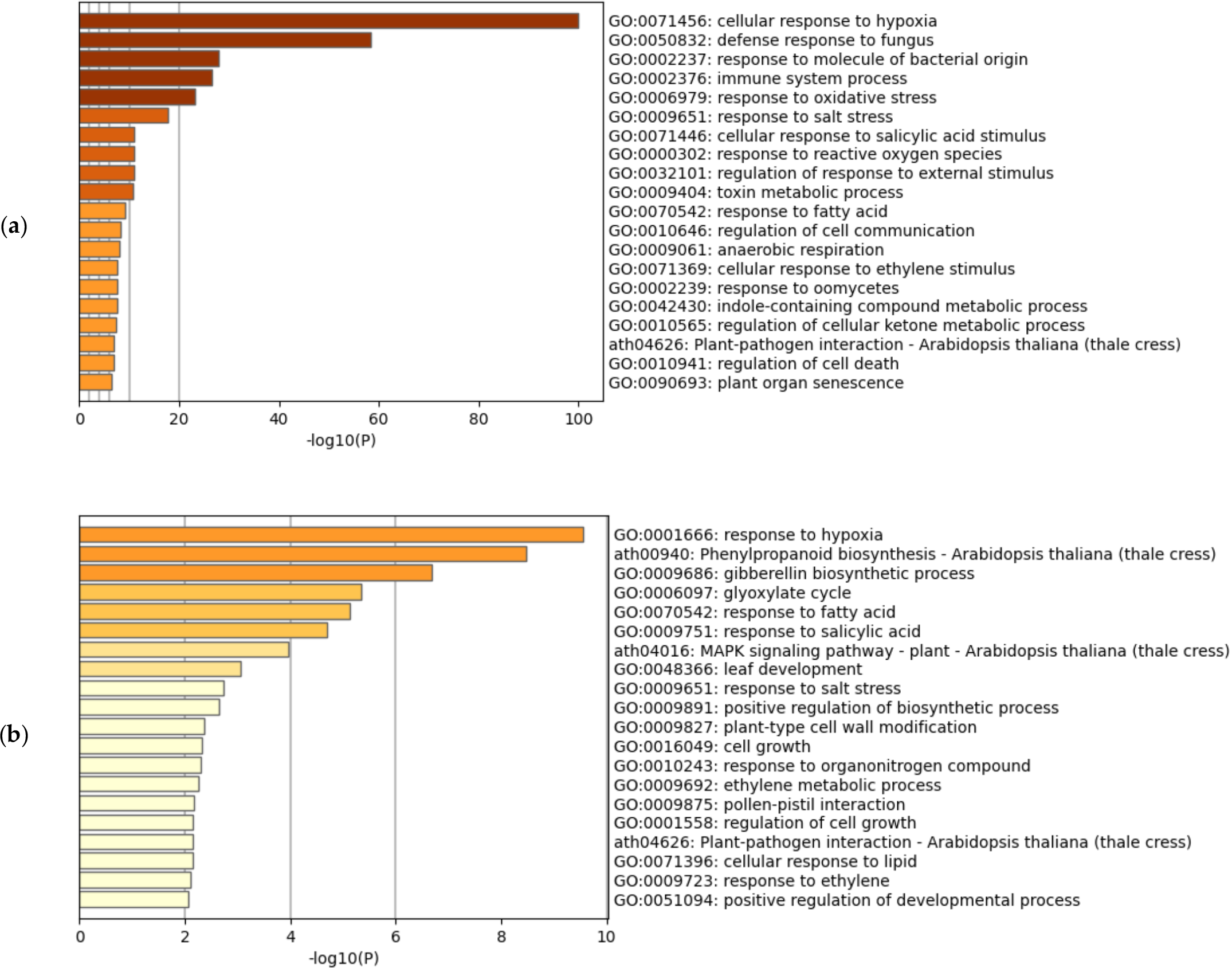
Gene set enrichment analysis of hypoxia-inducible upregulated genes. (**a**) Meta-analysis of Arabidopsis; (**b**) Meta-analysis of rice using the corresponding Arabidopsis genes.

The meta-analysis showed that the representative hypoxia-related genes in Arabidopsis were classified in accordance with previous studies (Table 2). Among the five Arabidopsis ERFVII TFs, the expression of *HRE1* and *HRE2* was upregulated, whereas *RAP2*.*12, RAP2*.*2*, and *RAP2*.*3* expression was unchanged (Table 2). All Arabidopsis core HRGs (*LBD41, PCO1, PCO2, ADH1*, and *PDC1*) regulated by *RAP2*.*2* and *RAP2*.*12* [11] were also upregulated (Table 2). PLANT CYSTEINE OXIDASEs (PCO1 and PCO2) are involved in the stability of ERFVII TFs [33], and ALCOHOL DEHYDROGENASE 1 (ADH1) and PYRUVATE DECARBOXYLASE 1 (PDC1) are involved in fermentative metabolism [6]. The expression of rice ERFVII TFs *SUB1B* and *SUB1C* was also upregulated (figshare file 5b) [19]. Considering that most of the experiments on rice involved the cultivar Nipponbare and that the Nipponbare transcriptome was used as a reference, we excluded submergence-tolerant ERFVII TFs (*SUB1A, SK1*, and *SK2*), which were absent in the Nipponbare genome [7,9], from our meta-analysis.

**Table 2.**
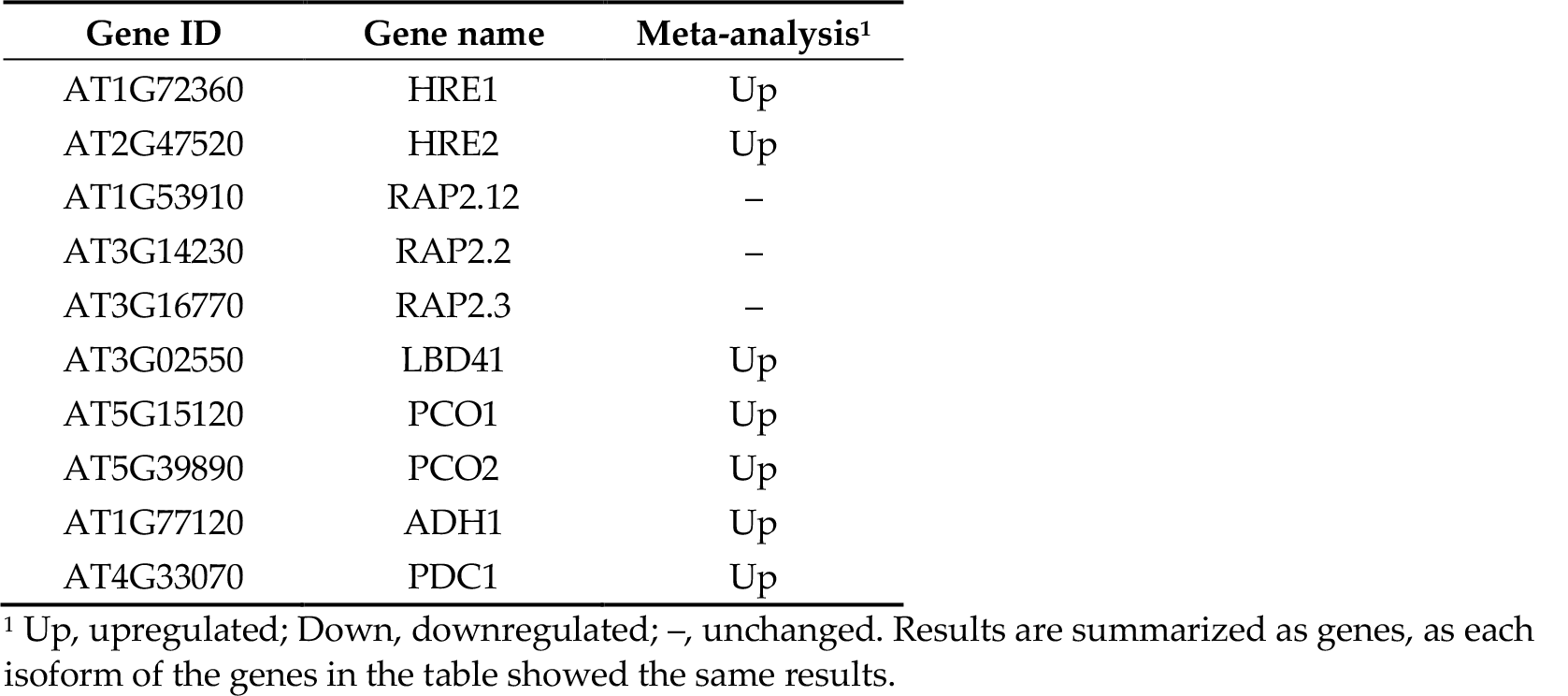
HN-scores of representative transcripts related to hypoxia response in Arabidopsis.

### 3.3. Characteristics of Downregulated Transcripts under Hypoxia

Gene set enrichment analysis of the downregulated transcripts revealed different trends between Arabidopsis and rice (Figure 2a,b). In Arabidopsis, the most significantly enriched term was GO:0009642 (response to light intensity) (Figure 2a). Darkness is often associated with natural submergence [6], some experiments were collected for the meta-analysis of Arabidopsis conducted under darkness and submergence (figshare file 1a) [19]. These conditions can downregulate genes related to response to light intensity. However, when we analyzed hypoxic treatments only, the most significantly enriched term was GO:0016049 (cell growth) (figshare file 7) [19], suggesting that the presence of water or darkness also affects change of gene expression. The lowest HN-score of −21 was observed in AT5G65730.1 (XTH6) (figshare file 5c) [19]. Some members of the XYLOGLUCAN ENDOTRANSGLUCOSYLASE/HYDROLASE (XTH) gene family have been studied in relation to cell wall expansion [34,35]. In rice, the most significantly enriched term was GO:0006790 (sulfur compound metabolic process) (Figure 2b). Ethylene decreases the antioxidant metallothionein, a small cysteine-rich protein [6,36]. The downregulated genes in the term GO:0006790 may be related to a decrease in metallothionein. The lowest HN-score of −22 was observed in Os05t0217700-01 (OsBURP07) and two isoforms of Os08g0531000 (Os08t0531000-01 and Os08t0531000-02) (figshare file 5d) [19]. OsBURP07 is downregulated under abscisic acid (ABA) treatment [37]; however, this phenomenon cannot be attributed to the ABA treatment because ethylene decreases ABA levels in submerged tissues [38]. Os08g0531000, also known as NUCLEOTIDE PYROPHOSPHATASE/PHOSPHODIESTERASE 1, negatively regulates starch accumulation and growth [39]. We also performed the gene set enrichment analysis against GO Molecular Function and GO Cellular Component (figshare file 9) [19]. In GO Molecular Function, GO:0046906 (tetrapyrrole binding) was commonly enriched in Arabidopsis and rice. This member includes many of the cytochrome P450 monooxygenases, which play important role in metabolic diversification [40]. The enrichment of this GO may be related to decreased activity of metabolic processes.

**Figure 2.**
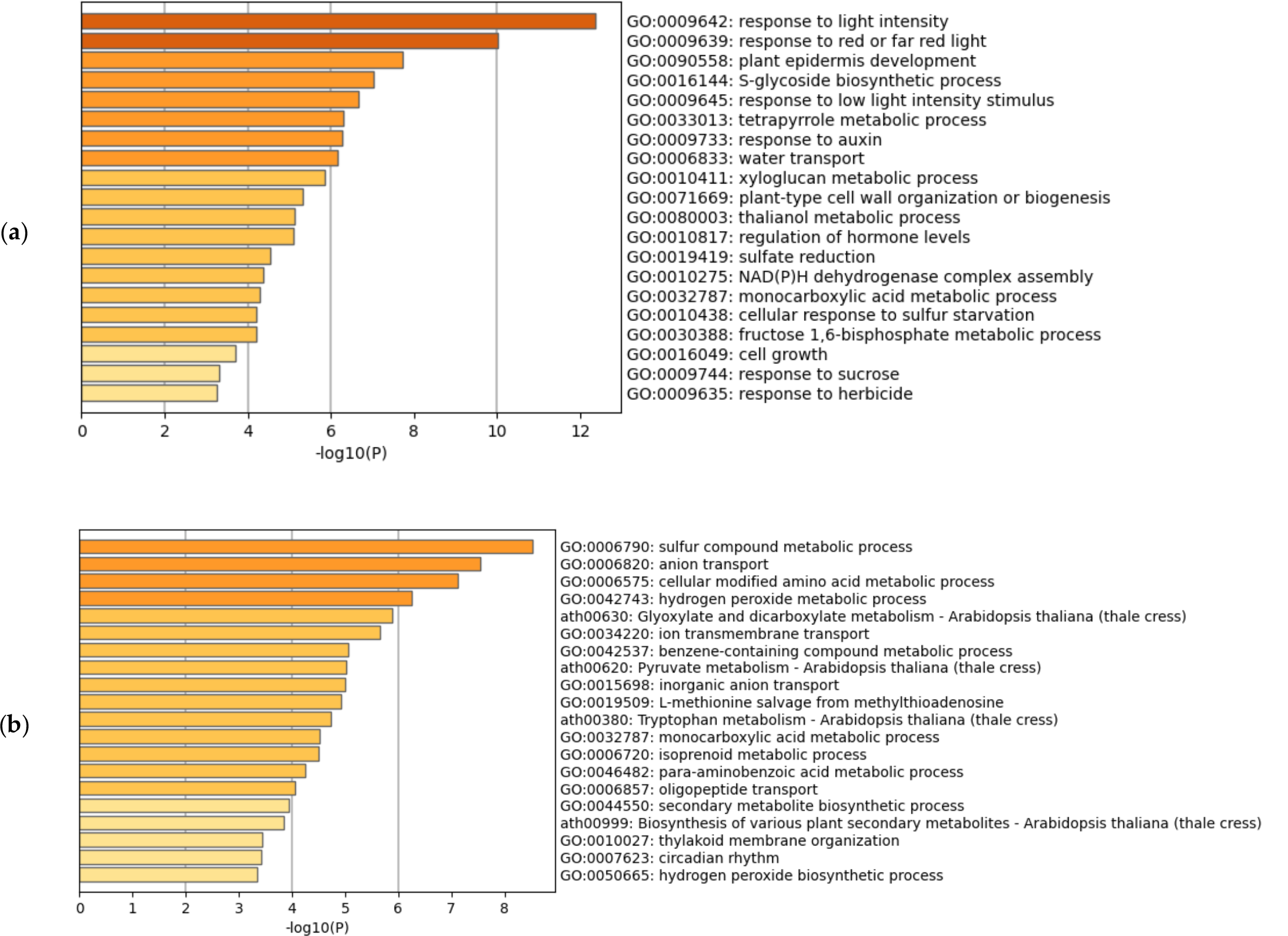
Gene set enrichment analysis of hypoxia-inducible downregulated genes. (**a**) Meta-analysis of Arabidopsis. (**b**) Meta-analysis of rice using the corresponding Arabidopsis genes.

### 3.4. Identification of Commonly Upregulated or Downregulated Genes in Arabidopsis and Rice

To identify novel candidate genes related to hypoxia in plants, we focused on the genes commonly upregulated or downregulated in Arabidopsis and rice because they are expected to be related to common mechanisms in plants. Upregulated or downregulated transcript IDs, converted to orthologs in Arabidopsis, were summarized as Arabidopsis gene IDs, and overlaps between the two species were visualized (Figure 3a,b). We identified 40 and 19 commonly upregulated and downregulated genes, respectively (Tables 3 and 4, respectively). Gene set enrichment analysis of the overlapping genes showed that the most significantly enriched term of the upregulated genes was GO:0071456 (cellular response to hypoxia), whereas the downregulated genes showed a less confident enriched term (Figure 3c,d).

**Figure 3.**
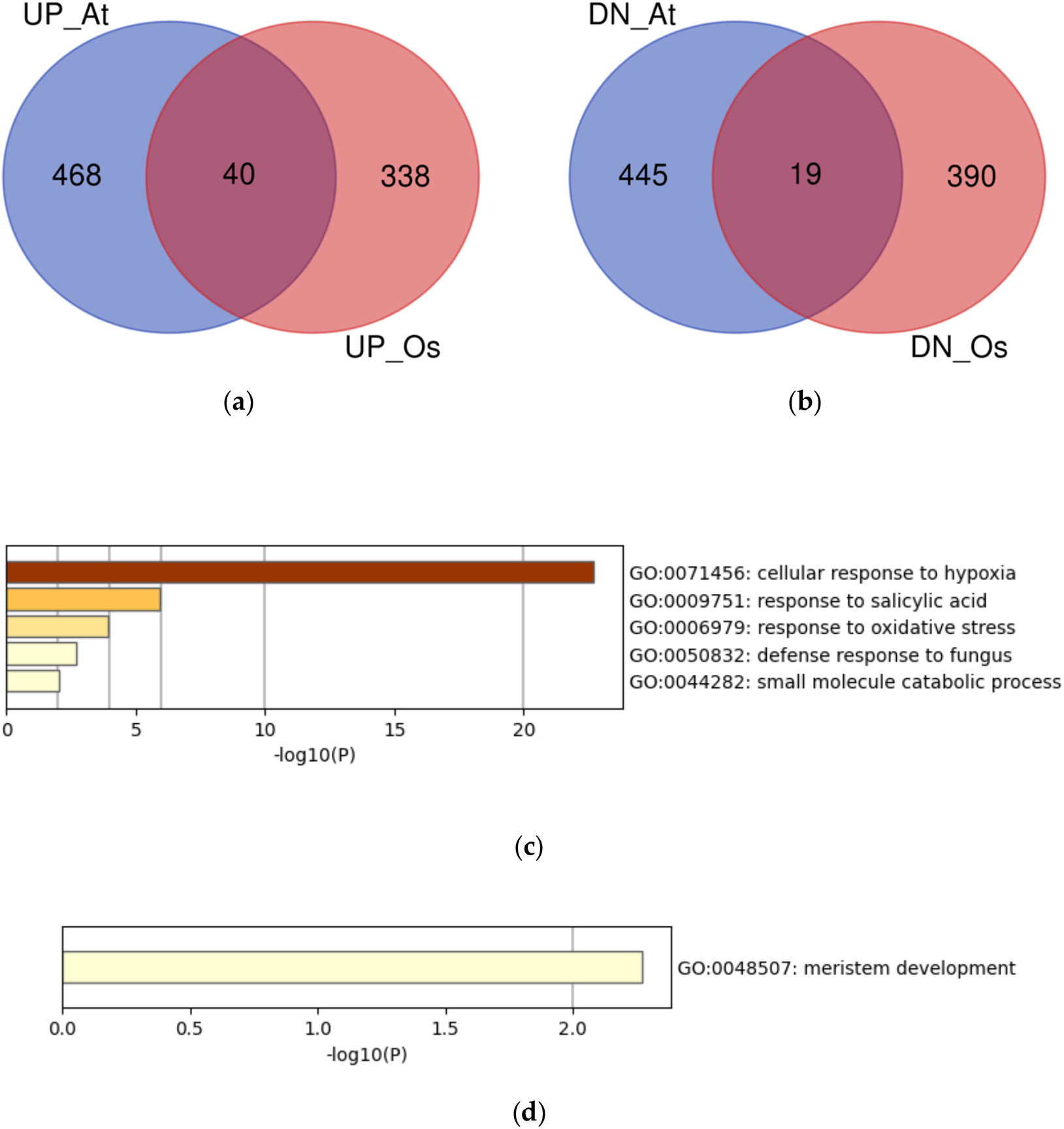
Analysis of the overlaps of upregulated or downregulated genes in Arabidopsis and rice using the Arabidopsis gene IDs. (**a**) Venn diagram of the upregulated genes in Arabidopsis (UP_At) and rice (UP_Os). (**b**) Venn diagram of the downregulated genes in Arabidopsis (DN_At) and rice (DN_Os). (**c**) Gene set enrichment analysis of the commonly upregulated genes. (**d**) Gene set enrichment analysis of the commonly downregulated genes.

**Table 3.** Commonly upregulated transcripts in Arabidopsis and rice, indicated as Arabidopsis gene IDs.

| Gene ID | Gene name | in<br>GO:0071456<sup>1</sup> | Gene id | Gene name | in<br>GO:0071456<sup>1</sup> |
| --- | --- | --- | --- | --- | --- |
| AT1G80840 | WRKY40 | – | AT2G46400 | WRKY46 | Yes |
| AT1G77120 | ADH1 | Yes | AT3G11820 | SYP121 | – |
| AT5G24530 | DMR6 | Yes | AT5G39890 | PCO2 | Yes |
| AT4G20860 | FAD-OXR | Yes | AT3G11930 | – | – |
| AT3G22060 | CRRSP38 | – | AT2G23810 | TET8 | – |
| AT1G18390 | – | – | AT4G10265 | – | Yes |
| AT1G15670 | – | – | AT5G06320 | NHL3 | Yes |
| AT1G72210 | BHLH96 | – | AT1G43800 | S-ACP-DES6 | Yes |
| AT3G61060 | AtPP2-A13 | – | AT2G38470 | WRKY33 | – |
| AT4G33050 | EDA39 | – | AT5G50200 | NRT3.1 | – |
| AT5G47060 | – | Yes | AT3G29970 | – | – |
| AT4G10270 | – | – | AT3G02550 | LBD41 | Yes |
| AT1G72360 | HRE1 | Yes | AT2G30490 | CYP73A5 | – |
| AT4G23810 | WRKY53 | – | AT4G27450 | – | Yes |
| AT2G47520 | ERF071 | Yes | AT1G68840 | RAV2 | Yes |
| AT1G14870 | PCR2 | – | AT2G19590 | ACO1 | – |
| AT5G41080 | GDPD2 | Yes | AT2G40000 | HSPRO2 | Yes |
| AT5G48540 | CRRSP55 | – | AT2G36690 | – | – |
| AT1G25560 | TEM1 | Yes | AT5G06570 | – | – |
| AT5G25930 | – | – | AT5G64120 | PER71 | – |

<sup>1</sup> GO:0071456 (cellular response to hypoxia)**Table 4.** Commonly downregulated transcripts in Arabidopsis and rice, indicated as Arabidopsis gene IDs.

**Table 4.** Commonly downregulated transcripts in Arabidopsis and rice, indicated as Arabidopsis gene IDs.

| Gene ID | Gene name | Gene ID | Gene name | Gene ID | Gene name |
| --- | --- | --- | --- | --- | --- |
| AT4G35100 | PIP2;7 | AT2G39890 | PROT1 | AT5G50740 | – |
| AT3G45780 | PHOT1 | AT2G29630 | THIC | AT5G57180 | CIA2 |
| AT2G38760 | ANN3 | AT1G28570 | – | AT4G23400 | PIP1;5 |
| AT4G18570 | – | AT1G16880 | ACR11 | AT3G55760 | – |
| AT4G19170 | CCD4 | AT5G42510 | DIR1 | AT1G62180 | APR2 |
| AT1G20020 | LFNR2 | AT3G18000 | NMT1 |  |  |
| AT5G11420 | – | AT5G05980 | FPGS1 |  |  |

Among the 40 commonly upregulated genes, 23 were not included in the term GO:0071456, suggesting that these may be novel genes upregulated by hypoxia (Table 3). Notably, three of the four WRKY TFs that were commonly upregulated were not included in the term GO:0071456. CYP73A5 is C4H, which is a key enzyme in the phenylpropanoid pathway [41]. *ACC OXIDASE 1* (*ACO1*) does not include the term GO:0071456, but upregulation of *ACO1* expression under hypoxia has been reported in Arabidopsis [42]. Among the 19 commonly downregulated genes were aquaporin genes *PLASMA MEMBRANE INTRINSIC PROTEIN 2;7* (*PIP2*;7) and *PIP1;5* (Table 4). PIP2;7 is an active water channel in Arabidopsis [43]; hence, the downregulation of aquaporin genes may be related to the inhibition of excess water transport under submergence. We also analyzed the possible protein–protein interactions of the commonly upregulated and downregulated genes using STRING (figshare file 10) [19]. Although most of the commonly upregulated genes were in association with co-expression and text mining evidence, none of the known or predicted interactions was observed.

## 4. Discussion

In this study, we collected 29 (Arabidopsis) and 26 (rice) pairs of RNA-Seq data of hypoxic and normoxic treatments and performed a meta-analysis of the changes in gene expression under hypoxia. We treated biological replicates from the same series of experiments as an individual experiment, and confirmed the swapping of the pairs in the replicates produced similar results in gene set enrichment analysis (figshare file 11) [19]. Although the present meta-analysis included fewer experiments than a similar meta-analysis performed in humans [15], it clearly showed the enrichment of hypoxia-related GO terms in Arabidopsis and rice (Figure 1a,b). This result suggests that the present meta-analysis reflects the global trends of differential gene expression under hypoxia. Moreover, the representative hypoxia-related genes in Arabidopsis showed the same expression patterns as those in previous studies (Table 2). We believe that the dataset used in our meta-analysis correctly reflects our scientific knowledge of gene expression under hypoxia.

Our meta-analysis aimed to elucidate the unknown molecular mechanisms. To this end, we focused on the commonly upregulated or downregulated genes in Arabidopsis and rice. We identified 40 upregulated and 19 downregulated genes in both species (Tables 3 and 4). Among the 40 commonly upregulated genes, four WRKY TFs were included, of which three were not included in the hypoxia-related GO terms (Table 3). The role of WRKY TFs in the hypoxia response has not been studied in Arabidopsis, but in persimmon (*Diospyros kaki*), the expression of some WRKY TFs, including *DkWRKY1*, is upregulated by hypoxia and the process involves *DkPDC2* transactivation [44]. Another report showed that rice *WRKY62* is upregulated under hypoxia and activates hypoxia genes, while repressing defense-related diterpenoid phytoalexin factor under hypoxia [45]. However, in the present meta-analysis, *WRKY62* (Os09t0417800-01 and Os09t0417800-02) expression was not upregulated. Determining the role of WRKY TFs, including the four commonly upregulated genes, in hypoxia response is a valuable research goal. We also detected an upregulation in expression of *CYP73A5* (*C4H*), a key enzyme of the phenylpropanoid pathway, in Arabidopsis and rice, and its corresponding gene was not included in the hypoxia-related GO term (Table 3). However, the upstream gene [phenylalanine ammonia-lyase (*PAL*)] and the downstream gene [4-coumaroyl CoA ligase (*4CL*)] of C4H in the general phenylpropanoid pathway were not upregulated in either Arabidopsis or rice (figshare files 6a,b) [19]. A previous study suggested that flavonoid and lignin biosyntheses are suppressed in Arabidopsis under hypoxia [31]. The effect of *C4H* upregulation under hypoxia may be the subject of future studies.

Finally, we compared our results with those of a previous meta-analysis of hypoxia in human cell lines and tissue specimens [15]. In our meta-analysis, two well-studied HRGs, namely, *ADH1* and *PCO2*, were upregulated in both Arabidopsis and rice (Table 3). The orthologs in humans identified in this study are ADH5 and ADO, respectively (figshare file 5a) [19]. A previous meta-analysis of hypoxia in humans obtained HN-scores (1.5-fold threshold; 495 experimental pairs) of −34 and −84 for ADH5 and ADO, respectively [46], indicating that their orthologs in humans are downregulated under hypoxia. The fermentative metabolism of pyruvate is directed toward lactate in animals via lactate dehydrogenase (LDH), in contrast to ethanol formation in plants via PDC and ADH [26]. Indeed, the HN-score of LDHA in a previous meta-analysis in humans was 334, wherein it ranked in the top 100 upregulated genes [47]. ADO is an enzymatic oxygen sensor with an equivalent role in plant PCOs, identified as a component of an alternative mechanism for oxygen-sensitive proteolysis in mammals [3,48]. Our meta-analysis suggests distinct molecular mechanisms under hypoxia in plants and animals.

## Author Contributions

Conceptualization, K.T. and H.B.; methodology, K.T. and H.B.; software, K.T.; validation, K.T. and H.B.; formal analysis, K.T.; investigation, K.T.; resources, H.B.; data curation, K.T.; writing—original draft preparation, K.T.; writing—review and editing, K.T. and H.B.; visualization, K.T.; supervision, H.B.; project administration, H.B.; funding acquisition, H.B. All authors have read and agreed to the published version of the manuscript.

## Funding

This work was supported by the Center of Innovation for Bio-Digital Transformation (BioDX), an open innovation platform for industry-academia co-creation of Japan Science and Technology Agency (JST, COI-NEXT, JPMJPF2010).

## Institutional Review Board Statement

Not applicable.

## Informed Consent Statement

Not applicable.

## Data Availability Statement

The data presented in this study are openly available in figshare [19].

## Acknowledgments

We would like to thank Dr. Kouhei Toga and Mr. Takayuki Suzuki for their valuable discussion.

## Conflicts of Interest

The authors declare no conflict of interest.

## Notes

https://doi.org/10.6084/m9.figshare.20055086

### Competing Interest Statement

The authors have declared no competing interest.

### Summary of Updates

Revision minor issues

## References

1. Kaur, G.; Singh, G.; Motavalli, P.P.; Nelson, K.A.; Orlowski, J.M.; Golden, B.R. Impacts and Management Strategies for Crop Production in Waterlogged or Flooded Soils: A Review. Agron. J. 2020, 112, 1475–1501, doi:10.1002/agj2.20093.

2. Fukao, T.; Barrera-Figueroa, B.E.; Juntawong, P.; Peña-Castro, J.M. Submergence and Waterlogging Stress in Plants: A Review Highlighting Research Opportunities and Understudied Aspects. Front. Plant Sci. 2019, 10, 340, doi:10.3389/fpls.2019.00340.

3. Holdsworth, M.J.; Gibbs, D.J. Comparative Biology of Oxygen Sensing in Plants and Animals. Curr. Biol. CB 2020, 30, R362–R369, doi:10.1016/j.cub.2020.03.021.

4. Doorly, C.M.; Graciet, E. Lessons from Comparison of Hypoxia Signaling in Plants and Mammals. Plants 2021, 10, 993, doi:10.3390/plants10050993.

5. Kaelin, W.G.; Ratcliffe, P.J. Oxygen Sensing by Metazoans: The Central Role of the HIF Hydroxylase Pathway. Mol. Cell 2008, 30, 393–402, doi:10.1016/j.molcel.2008.04.009.

6. Loreti, E.; van Veen, H.; Perata, P. Plant Responses to Flooding Stress. Curr. Opin. Plant Biol. 2016, 33, 64–71, doi:10.1016/j.pbi.2016.06.005.

7. Xu, K.; Xu, X.; Fukao, T.; Canlas, P.; Maghirang-Rodriguez, R.; Heuer, S.; Ismail, A.M.; Bailey-Serres, J.; Ronald, P.C.; Mackill, D.J. Sub1A Is an Ethylene-Response-Factor-like Gene That Confers Submergence Tolerance to Rice. Nature 2006, 442, 705–708, doi:10.1038/nature04920.

8. Fukao, T.; Xu, K.; Ronald, P.C.; Bailey-Serres, J. A Variable Cluster of Ethylene Response Factor–Like Genes Regulates Metabolic and Developmental Acclimation Responses to Submergence in Rice. Plant Cell 2006, 18, 2021–2034, doi:10.1105/tpc.106.043000.

9. Hattori, Y.; Nagai, K.; Furukawa, S.; Song, X.-J.; Kawano, R.; Sakakibara, H.; Wu, J.; Matsumoto, T.; Yoshimura, A.; Kitano, H.; et al. The Ethylene Response Factors SNORKEL1 and SNORKEL2 Allow Rice to Adapt to Deep Water. Nature 2009, 460, 1026–1030, doi:10.1038/nature08258.

10. Gibbs, D.J.; Conde, J.V.; Berckhan, S.; Prasad, G.; Mendiondo, G.M.; Holdsworth, M.J. Group VII Ethylene Response Factors Coordinate Oxygen and Nitric Oxide Signal Transduction and Stress Responses in Plants. Plant Physiol. 2015, 169, 23–31, doi:10.1104/pp.15.00338.

11. Gasch, P.; Fundinger, M.; Müller, J.T.; Lee, T.; Bailey-Serres, J.; Mustroph, A. Redundant ERF-VII Transcription Factors Bind to an Evolutionarily Conserved Cis-Motif to Regulate Hypoxia-Responsive Gene Expression in Arabidopsis. Plant Cell 2016, 28, 160–180, doi:10.1105/tpc.15.00866.

12. Licausi, F.; van Dongen, J.T.; Giuntoli, B.; Novi, G.; Santaniello, A.; Geigenberger, P.; Perata, P. HRE1 and HRE2, Two Hypoxia-Inducible Ethylene Response Factors, Affect Anaerobic Responses in Arabidopsis thaliana. Plant J. 2010, 62, 302–315, doi:10.1111/j.1365-313X.2010.04149.x.

13. Mustroph, A.; Zanetti, M.E.; Jang, C.J.H.; Holtan, H.E.; Repetti, P.P.; Galbraith, D.W.; Girke, T.; Bailey-Serres, J. Profiling Translatomes of Discrete Cell Populations Resolves Altered Cellular Priorities during Hypoxia in Arabidopsis. Proc. Natl. Acad. Sci. U. S. A. 2009, 106, 18843–18848, doi:10.1073/pnas.0906131106.

14. Gibbs, D.J.; Lee, S.C.; Md Isa, N.; Gramuglia, S.; Fukao, T.; Bassel, G.W.; Correia, C.S.; Corbineau, F.; Theodoulou, F.L.; Bailey-Serres, J.; et al. Homeostatic Response to Hypoxia Is Regulated by the N-End Rule Pathway in Plants. Nature 2011, 479, 415–418, doi:10.1038/nature10534.

15. Ono, Y.; Bono, H. Multi-Omic Meta-Analysis of Transcriptomes and the Bibliome Uncovers Novel Hypoxia-Inducible Genes. Biomedicines 2021, 9, 582, doi:10.3390/biomedicines9050582.

16. Bono, H.; Hirota, K. Meta-Analysis of Hypoxic Transcriptomes from Public Databases. Biomedicines 2020, 8, 10, doi:10.3390/biomedicines8010010.

17. Bono, H. All of Gene Expression (AOE): An Integrated Index for Public Gene Expression Databases. PLoS One 2020, 15, e0227076, doi:10.1371/journal.pone.0227076.

18. Barrett, T.; Wilhite, S.E.; Ledoux, P.; Evangelista, C.; Kim, I.F.; Tomashevsky, M.; Marshall, K.A.; Phillippy, K.H.; Sherman, P.M.; Holko, M.; et al. NCBI GEO: Archive for Functional Genomics Data Sets—Update. Nucleic Acids Res. 2013, 41, D991–D995, doi:10.1093/nar/gks1193.

19. Tamura, K. Meta-Analysis of RNA Sequencing Data of Arabidopsis and Rice under Hypoxia. figshare 2022, doi:10.6084/m9.figshare.20055086.

20. Patro, R.; Duggal, G.; Love, M.I.; Irizarry, R.A.; Kingsford, C. Salmon Provides Fast and Bias-Aware Quantification of Transcript Expression. Nat. Methods 2017, 14, 417–419, doi:10.1038/nmeth.4197.

21. Yates, A.D.; Allen, J.; Amode, R.M.; Azov, A.G.; Barba, M.; Becerra, A.; Bhai, J.; Campbell, L.I.; Carbajo Martinez, M.; Chakiachvili, M.; et al. Ensembl Genomes 2022: An Expanding Genome Resource for Non-Vertebrates. Nucleic Acids Res. 2022, 50, D996–D1003, doi:10.1093/nar/gkab1007.

22. Bono, H. Meta-Analysis of Oxidative Transcriptomes in Insects. Antioxidants 2021, 10, 345, doi:10.3390/antiox10030345.

23. Kinsella, R.J.; Kähäri, A.; Haider, S.; Zamora, J.; Proctor, G.; Spudich, G.; Almeida-King, J.; Staines, D.; Derwent, P.; Kerhornou, A.; et al. Ensembl BioMarts: A Hub for Data Retrieval across Taxonomic Space. Database 2011, bar030, doi:10.1093/database/bar030.

24. Zhou, Y.; Zhou, B.; Pache, L.; Chang, M.; Khodabakhshi, A.H.; Tanaseichuk, O.; Benner, C.; Chanda, S.K. Metascape Provides a Biologist-Oriented Resource for the Analysis of Systems-Level Datasets. Nat. Commun. 2019, 10, 1523, doi:10.1038/s41467-019-09234-6.

25. Szklarczyk, D.; Gable, A.L.; Lyon, D.; Junge, A.; Wyder, S.; Huerta-Cepas, J.; Simonovic, M.; Doncheva, N.T.; Morris, J.H.; Bork, P.; et al. STRING V11: Protein–Protein Association Networks with Increased Coverage, Supporting Functional Discovery in Genome-Wide Experimental Datasets. Nucleic Acids Res. 2019, 47, D607–D613, doi:10.1093/nar/gky1131.

26. Chakraborty, K.; Ray, S.; Vijayan, J.; Molla, K.A.; Nagar, R.; Jena, P.; Mondal, S.; Panda, B.B.; Shaw, B.P.; Swain, P.; et al. Preformed Aerenchyma Determines the Differential Tolerance Response under Partial Submergence Imposed by Fresh and Saline Water Flooding in Rice. Physiol. Plant. 2021, 173, 1597–1615, doi:10.1111/ppl.13536.

27. Mustroph, A.; Lee, S.C.; Oosumi, T.; Zanetti, M.E.; Yang, H.; Ma, K.; Yaghoubi-Masihi, A.; Fukao, T.; Bailey-Serres, J. Cross-Kingdom Comparison of Transcriptomic Adjustments to Low-Oxygen Stress Highlights Conserved and Plant-Specific Responses. Plant Physiol. 2010, 152, 1484–1500, doi:10.1104/pp.109.151845.

28. Yang, C.-Y.; Hong, C.-P. The NADPH Oxidase Rboh D Is Involved in Primary Hypoxia Signalling and Modulates Expression of Hypoxia-Inducible Genes under Hypoxic Stress. Environ. Exp. Bot. 2015, 115, 63–72, doi:10.1016/j.envexpbot.2015.02.008.

29. Kadota, Y.; Shirasu, K.; Zipfel, C. Regulation of the NADPH Oxidase RBOHD During Plant Immunity. Plant Cell Physiol. 2015, 56, 1472–1480, doi:10.1093/pcp/pcv063.

30. Lokdarshi, A.; Conner, W.C.; McClintock, C.; Li, T.; Roberts, D.M. Arabidopsis CML38, a Calcium Sensor That Localizes to Ribonucleoprotein Complexes under Hypoxia Stress. Plant Physiol. 2016, 170, 1046–1059, doi:10.1104/pp.15.01407.

31. Takahashi, H.; Yamauchi, T.; Rajhi, I.; Nishizawa, N.K.; Nakazono, M. Transcript Profiles in Cortical Cells of Maize Primary Root during Ethylene-Induced Lysigenous Aerenchyma Formation under Aerobic Conditions. Ann. Bot. 2015, 115, 879–894, doi:10.1093/aob/mcv018.

32. Liu, F.; VanToai, T.; Moy, L.P.; Bock, G.; Linford, L.D.; Quackenbush, J. Global Transcription Profiling Reveals Comprehensive Insights into Hypoxic Response in Arabidopsis. Plant Physiol. 2005, 137, 1115–1129, doi:10.1104/pp.104.055475.

33. Weits, D.A.; Giuntoli, B.; Kosmacz, M.; Parlanti, S.; Hubberten, H.-M.; Riegler, H.; Hoefgen, R.; Perata, P.; van Dongen, J.T.; Licausi, F. Plant Cysteine Oxidases Control the Oxygen-Dependent Branch of the N-End-Rule Pathway. Nat. Commun. 2014, 5, 3425, doi:10.1038/ncomms4425.

34. Kaewthai, N.; Gendre, D.; Eklöf, J.M.; Ibatullin, F.M.; Ezcurra, I.; Bhalerao, R.P.; Brumer, H. Group III-A XTH Genes of Arabidopsis Encode Predominant Xyloglucan Endohydrolases That Are Dispensable for Normal Growth. Plant Physiol. 2013, 161, 440–454, doi:10.1104/pp.112.207308.

35. Maris, A.; Suslov, D.; Fry, S.C.; Verbelen, J.-P.; Vissenberg, K. Enzymic Characterization of Two Recombinant Xyloglucan Endotransglucosylase/Hydrolase (XTH) Proteins of Arabidopsis and Their Effect on Root Growth and Cell Wall Extension. J. Exp. Bot. 2009, 60, 3959–3972, doi:10.1093/jxb/erp229.

36. Yamauchi, T.; Fukazawa, A.; Nakazono, M. METALLOTHIONEIN Genes Encoding ROS Scavenging Enzymes Are Down-Regulated in the Root Cortex during Inducible Aerenchyma Formation in Rice. Plant Signal. Behav. 2017, 12, e1388976, doi:10.1080/15592324.2017.1388976.

37. Ding, X.; Hou, X.; Xie, K.; Xiong, L. Genome-Wide Identification of BURP Domain-Containing Genes in Rice Reveals a Gene Family with Diverse Structures and Responses to Abiotic Stresses. Planta 2009, 230, 149–163, doi:10.1007/s00425-009-0929-z.

38. Fukao, T.; Bailey-Serres, J. Ethylene—A Key Regulator of Submergence Responses in Rice. Plant Sci. 2008, 175, 43–51, doi:10.1016/j.plantsci.2007.12.002.

39. Kaneko, K.; Inomata, T.; Masui, T.; Koshu, T.; Umezawa, Y.; Itoh, K.; Pozueta-Romero, J.; Mitsui, T. Nucleotide Pyrophosphatase/Phosphodiesterase 1 Exerts a Negative Effect on Starch Accumulation and Growth in Rice Seedlings under High Temperature and CO2 Concentration Conditions. Plant Cell Physiol. 2014, 55, 320–332, doi:10.1093/pcp/pct139.

40. Hansen, C.C.; Nelson, D.R.; Møller, B.L.; Werck-Reichhart, D. Plant Cytochrome P450 Plasticity and Evolution. Mol. Plant 2021, 14, 1244–1265, doi:10.1016/j.molp.2021.06.028.

41. Schilmiller, A.L.; Stout, J.; Weng, J.-K.; Humphreys, J.; Ruegger, M.O.; Chapple, C. Mutations in the Cinnamate 4-Hydroxylase Gene Impact Metabolism, Growth and Development in Arabidopsis. Plant J. 2009, 60, 771–782, doi:10.1111/j.1365-313X.2009.03996.x.

42. Ventura, I.; Brunello, L.; Iacopino, S.; Valeri, M.C.; Novi, G.; Dornbusch, T.; Perata, P.; Loreti, E. Arabidopsis Phenotyping Reveals the Importance of Alcohol Dehydrogenase and Pyruvate Decarboxylase for Aerobic Plant Growth. Sci. Rep. 2020, 10, 16669, doi:10.1038/s41598-020-73704-x.

43. Hachez, C.; Laloux, T.; Reinhardt, H.; Cavez, D.; Degand, H.; Grefen, C.; De Rycke, R.; Inzé, D.; Blatt, M.R.; Russinova, E.; et al. Arabidopsis SNAREs SYP61 and SYP121 Coordinate the Trafficking of Plasma Membrane Aquaporin PIP2;7 to Modulate the Cell Membrane Water Permeability. Plant Cell 2014, 26, 3132–3147, doi:10.1105/tpc.114.127159.

44. Zhu, Q.; Gong, Z.; Huang, J.; Grierson, D.; Chen, K.; Yin, X. High-CO 2 /Hypoxia-Responsive Transcription Factors DkERF24 and DkWRKY1 Interact and Activate DkPDC2 Promoter. Plant Physiol. 2019, 180, 621–633, doi:10.1104/pp.18.01552.

45. Fukushima, S.; Mori, M.; Sugano, S.; Takatsuji, H. Transcription Factor WRKY62 Plays a Role in Pathogen Defense and Hypoxia-Responsive Gene Expression in Rice. Plant Cell Physiol. 2016, 57, 2541–2551, doi:10.1093/pcp/pcw185.

46. Ono, Y. Score Based on the Ratio of Gene Expression between Hypoxic and Normoxic Conditions(HN-Score). figshare 2021, doi:10.6084/m9.figshare.14141135.v1.

47. Ono, Y. Genelist_Top 100 Human Genes Up-Regulated under Hypoxic Conditions. figshare 2021, doi:10.6084/m9.figshare.14141015.v1.

48. Masson, N.; Keeley, T.P.; Giuntoli, B.; White, M.D.; Puerta, M.L.; Perata, P.; Hopkinson, R.J.; Flashman, E.; Licausi, F.; Ratcliffe, P.J. Conserved N-Terminal Cysteine Dioxygenases Transduce Responses to Hypoxia in Animals and Plants. Science 2019, 365, 65–69, doi:10.1126/science.aaw0112.

